# Gene expression is globally regulated by interacting nucleobase supply and mRNA composition demand: a mechanism disrupted by multiple disease states and drug treatments

**DOI:** 10.1101/2024.12.23.630131

**Authors:** Benjamin S. Pickard

**Affiliations:** Strathclyde Institute of Pharmacy and Biomedical Sciences, University of Strathclyde, Glasgow, UK

## Abstract

Conventional expression studies quantify messenger RNA (mRNA) transcript levels gene-by-gene. We recently showed that protein expression is modulated at a global scale by amino acid availability, suggesting that mRNA expression levels might be similarly affected by nucleobase supply. Re-analysis of transcriptomic datasets confirmed that nucleobase supply and mRNA A+U:C+G sequence composition interact to shape a global profile of expression which can be represented by simple numerical outputs.

In mammals, each separate organ and cell-type displays a distinct baseline profile of expression, influenced by differentiation state. Expression profiles shift dynamically across the circadian day and the menstrual cycle. They are also significantly distorted by viral infection, multiple complex genetic disorders (including Alzheimer’s disease, schizophrenia, and autoimmune disorders), and after treatment with 115 of the 597 chemical entities analysed. These entities included known toxins, but also many commonly prescribed medications such as antibiotics and proton pump inhibitors, thus revealing a new mechanism of drug action and side-effect.

A role for nucleobase supply is supported by the actions of nucleobase analogue treatments and by a model of the nucleobase metabolism disorder, Lesch-Nyhan syndrome. On the demand-side, mRNAs at compositional extremes are over-represented in key gene ontologies including transcription and cell division, making these processes particularly sensitive to swings in global expression. This permits efficient *en bloc* reprogramming of cell state through simple changes in nucleobase proportion and supply. It is also proposed that this mechanism helped mitigate the loss of essential amino acid synthesis in higher organisms.

In summary, global expression regulation is invisible to conventional transcriptomic analysis, but its measurement allows a useful distinction between active, promoter-mediated gene expression changes and passive, cell state-dependent transcriptional competence. Linking metabolism directly to expression offers an entirely new perspective on evolution, disease aetiopathology (including GxE interactions), and the nature of the pharmacological response.

## Introduction

To orchestrate complex programmes of embryonic development and adult organism homeostasis, static chromosomal DNA must generate highly dynamic, promoter-regulated profiles of messenger RNA (mRNA) and, subsequently, functional protein expression. In contrast to the documented complexities of promoter regulation, the actual molecular syntheses of mRNA and protein are largely ignored as a straightforward and frictionless transfer of encoded information. However, both processes are vast enzyme-mediated polymerisation reactions taking place within partially sealed vessels (the nucleus, the cell) in which substrates (here, nucleobases or amino acids) are finite. Substrate concentration is a key determinant of biochemical reaction rate and, therefore, warrants consideration as a potential regulator of expression.

We recently reported that situational shortages in amino acid supply from diet or biosynthesis interact with the amino acid composition demands of each protein to modulate global protein expression levels, agnostic of mRNA abundance [1]. In essence, the global protein expression we described can be thought of as the ‘wood’ to the ‘trees’ of individual protein expression, offering a high-level diagnostic of cell state. Here, the natural follow-on question is posed: does a reduced availability of RNA nucleobases (for example through disease or drug treatment) influence global gene expression levels independently of promoter activity? Answering this question for global protein expression required the stratification of proteomic expression data by the relative composition of the three nutritional subgroups of amino acids (essential amino acids, EAA, required from diet; non-essential amino acids, NEAA, synthesised within the body; and conditionally essential amino acids, CEAA, inefficiently synthesised) within each protein. This was a logical choice because the subgroups reflect dietary/biosynthetic constraints that could drive evolutionary adaptation. For mRNA expression, an analogous constraint would be insufficient supply of purine (A/G) or pyrimidine (C/U) nucleobases through altered biochemical synthesis or salvage [2]. However, the prevailing view has always been that nucleobase supply is tightly and co-ordinately regulated to preserve availability for DNA and RNA synthesis [3]. Nevertheless, a re-examination of 45 publicly available transcriptomic datasets was carried out using a new approach which assesses the influence of nucleobase composition on the expression level of each mRNA sequence. Using this approach, a simple representation of the global gene expression profile can be determined and compared across different tissues, disease states, and drug treatments.

## Results

### A global level of gene expression control is detectable after stratification of transcriptomic data by mRNA nucleobase composition

Gene transcript levels were examined at a global level in multiple publicly available gene expression datasets. **Fig. 1** outlines the concept and process used here to achieve this goal and to visualise outcomes. The typical high dynamic range (2^3^ to 2^15^) of individual gene expression is illustrated for over 30,000 transcripts detected by microarray in the rat liver [4] when treated with water (gavage) as control (**Fig.1a**). **Fig.1b** shows analysis of the same data, except average transcript expression was calculated and plotted within 25 equal-sized bins each containing hundreds of randomly allocated transcripts. This generates a simplified profile of expression for the control and in livers from rats treated with example agents, N-nitrosodimethylamine and glipizide. As expected, the plot is largely flat. However, when the same transcripts were ranked and binned along the x-axis according to increasing C nucleobase composition (**Fig.1c**), substantial influence on average bin expression level becomes apparent. A similar approach with A nucleobase composition also shows correlation with average expression level (**Fig.1d**).

**Figure 1.**
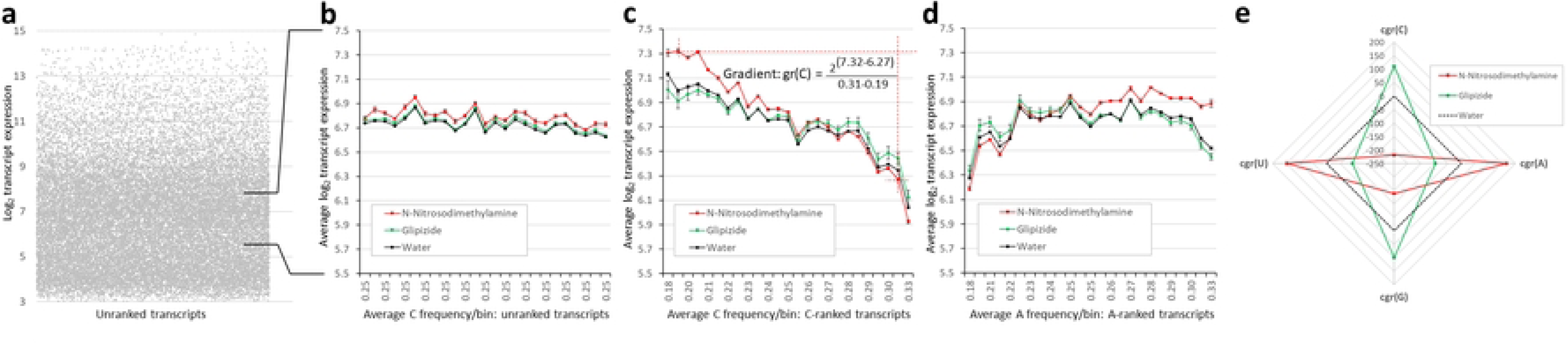
Stratification of expression data by mRNA base composition reveals transcript level regulation at a global scale. Rat liver gene expression changes (GEO dataset GSE57815 [4]) after in vivo treatment with N-Nitrosodimethylamine or Glipizide are used here to illustrate quantification of global expression profile. The complexity of individual transcript expression (a) can be removed by binning transcripts and plotting each bin’s average gene expression (b). A higher expression baseline after N-Nitrosodimethylamine treatment may represent a failure to normalise the dataset before public deposition of data. Both intrinsic and drug-induced global effects on expression, likely the result of limiting nucleobase availability, become apparent when transcripts are binned according to increases in their ranked composition, for example by nucleobase C (c) or A (d), then expression averaged per bin, and plotted (with error bars: s.e.m., N=3). In (c), the gradient of expression change (gr(C)) for N-Nitrosodimethylamine is shown calculated between the 2^nd^ to the 24^th^ C bin, chosen to avoid extreme outlier effects. By subtracting the control gradient, a corrected gradient, cgr(base), can be calculated for both drugs. A radar plot allows drug effects on cgr to be viewed simultaneously for all bases (e).

There is an intrinsic mRNA nucleobase composition effect on expression visible in control samples: a move from lowest to highest C composition bin is accompanied by a halving of average expression level. For A, it is more complex with a general positive influence on expression except at the extremes of composition. N-nitrosodimethylamine and glipizide treatments further distort global expression assessed by composition, but in opposing directions, resulting in anticlockwise and clockwise gradient changes, respectively, when ranking by A. To obtain numerical values for the action of a drug (or disease, see below) on expression, the gradient of expression change was calculated, as shown in **Fig.1c** for nucleobase C: **gr(C)**. The gradient for water, or appropriate solvent control, can be subtracted from this value to give a corrected value, **cgr(C)**, that offers some independence from microarray chip-specific effects or undocumented expression normalisation procedures, thus permitting improved cross-study comparison. It is hypothesised that changing gradients indicate the impact of a drug on nucleobase availability for incorporation into growing transcripts. Here, N-nitrosodimethylamine appears to be reducing C availability, further increasing constraint on transcripts with higher proportions of C and, thus, decreasing their expression level. Those transcripts with low levels of C show *relatively* enhanced expression when treated with N-nitrosodimethylamine. In **Fig.1e**, corrected gradient values for the two drugs and water control are shown on a radar plot, allowing drug influences on all four mRNA nucleobases to be visualised simultaneously, with the control gradient represented by the 0 contour (dashed line). This form of plot highlighted an unexpected finding - that drug-induced perturbations in gene expression profile do not vary in dimensions aligned with a simple purine (gr(A), gr(G)) or pyrimidine (gr(C), gr(U)) biochemical pathway disruption. Instead, global gene expression distortions primarily move along gr(C) correlated with gr(G), and gr(A) correlated with gr(U) directions. Henceforth, gr(C) and gr(A) alone are plotted as representatives of these two dimensions, and any shift in transcriptional profile is termed as being in either the ‘C-A+’ or ‘C+A-’ direction.

### Nucleobase composition differences between mRNA subregions underpin global transcriptional control and cause unexpected coding biases in proteins

The subregional features of mRNA composition were examined as a potential source of the A+U and C+G pairings of gradient movement. Within 110,962 human mRNA transcripts (protein-coding component of the GENCODE v44), the average nucleobase frequencies across the human transcriptome are very similar: 0.249 (C), 0.257 (A), 0.256 (G), and 0.238 (U) (Fig. 2a) and average transcript length is 2,395 nt. However, analysis of mRNA subregions revealed distinct base profiles: the 5’ Untranslated Region (UTR) shows enrichment for C and G bases (Fig.2b), likely correlating with promoter CpG island proximity; the protein-coding sequence (CDS), constrained by its coding function, shows no clear bias (Fig.2c); while the 3’ UTR shows enrichment for A and U bases (Fig.2d). Shorter mRNAs are disproportionately (C+G) rich on average (Fig.2e), but longer ones, by virtue of their proportionally greater CDS and 3’UTR contribution to length (Fig.2f), tend to greater (A+U) richness. These pairings of mRNA nucleobase composition frequencies match the nucleobase pairings observed for the dimensions of global transcriptional change. This suggests a model whereby nucleobase availability (supply) may be variable in nature and direction, but transcript composition (demand) dictates its consequences on expression, restricting global profile changes to the C+A- or C-A+ type.

**Figure 2.**
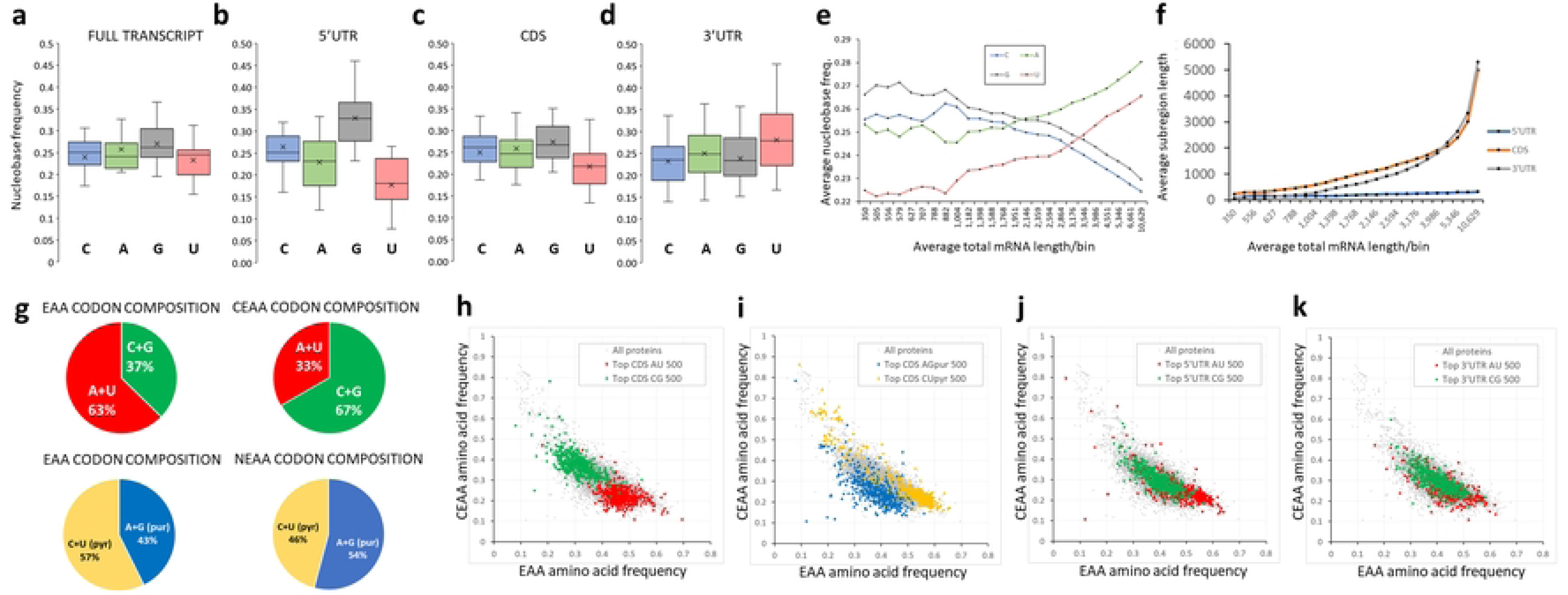
Distinct A+U and C+G base frequencies within subregions of mRNA transcripts are responsible for the bidirectionality of global expression changes and the biases in resulting protein composition. The proportions of the four nucleobases were quantified in all human full-length mRNA transcripts (a) and then by subregion: 5’UTR sequences (b), CDS coding sequences (c), and 3’UTR sequences (d) with results displayed as box-and-whisker plots. The differences in nucleobase composition between subregions show correlations between C and G (both decreasing in frequency) and, separately, between A and U bases (increasing) as the length of transcript increases (e). This length effect in nucleobase representation is explained by total transcript length increases deriving primarily from increases in A+U enriched CDS and 3’UTR sequences (f). In (g), the sets of codons for the nutritional groupings of amino acids show biases in nucleobase frequency. EAA: Essential amino acids; CEAA: conditionally essential amino acids; NEAA: non-essential amino acids. The consequences of these biases are revealed in plots of all proteins (h-k). The top 500 mRNAs for enrichment of A+U or C+G nucleobases code for proteins enriched in EAAs or CEAAs, respectively (h). A similar but weaker effect is seen for extreme purine- and pyrimidine-rich mRNAs and EAAs and NEAAs in their proteins, respectively, with NEAA-encoding indirectly shown by low EAA/CEAA values (i). mRNA-protein composition correlations are not only driven by direct action of CDS sequences – extreme A+U composition biases in the 5’UTR sequences (j) and 3’UTR sequences (k) also influence EAA/CEAA coding biases in proteins, indicating that detection of nucleobase supply (and impact on transcriptional competence) is transcript-wide.

From first principles, CDS sequences with extreme nucleobase compositions would be expected to encode proteins with a biased amino acid profile. However, the precise nature of this skew was unexpected. A+U and C+G pairings have very different relative representations within the codons for the 9 essential amino acids (EAA) compared to the codons for the 6 conditionally essential (CEAA) amino acids (**Fig.2g top**). To a lesser extent this is also true for the comparison of C+U (pyrimidine) and A+G (purine) pairings, which showed representational differences between EAA codons and those for the 5 non-essential amino acids (NEAA) (**Fig.2g bottom**). The consequence of a connection between nucleobase composition and nutritional amino acid groupings is that mRNAs of extreme composition (e.g., the 500 most A+U-rich or C+G-rich in the CDS subregion) encode highly EAA- or CEAA-enriched proteins (p-value comparing EAA composition between the 500 top A+U nucleobase enriched proteins against all proteins: 1.1 x 10^-69^, p-value comparing CEAA composition between the 500 top C+G enriched proteins against all proteins: 8.0 x 10^-100^) (**Fig.2h**). Purine/pyrimidine-rich CDS sequences showed less of a coding discrimination between EAA and NEAA groupings (**Fig.2i**).

Equally unexpectedly, the A+U influence on amino acid composition was not limited to the direct coding effects of the CDS region. Compositionally extreme non-coding 5’UTR or 3’UTR sequences, considered in isolation, also showed a significant influence on associated protein amino acid compositions (5’UTR effect p-values: EAA 2.1 x 10^-64^, CEAA 1.2 x 10^-27^; 3’UTR effect p-values: EAA 2.8 x 10^-17^, CEAA 1.7 x 10^-19^) (**Fig.2j,k)**. This influence was further confirmed by the modest positive correlations observed between sequence A+U composition frequencies within the isolated CDS sequences and the adjoining 5’UTR (R^2^ 0.12) or 3’UTR (R^2^ 0.15) sequences. This phenomenon did not occur with the purine/pyrimidine pairings.

To summarise this section, nucleobase composition variation is largely limited to the A+U and C+G dimensions by a mRNA subregion interaction with length, and this has undergone evolutionary selection along the *entire* length of mRNA transcripts to permit heightened sensitivity to nucleobase supply in a manner that affects both global mRNA expression as well as the nutritional/biosynthetic amino acid composition of encoded proteins.

### Global gene expression profiles are perturbed by many drugs/chemical entities

Thirteen publicly available drug-profiling gene expression datasets from human, mouse and rat (Gene Expression Ominibus, [5, 6]) were chosen for analysis, comprising induced pluripotent cell lines (partially differentiated down neural progenitor or cardiomyogenic lineages), transformed cell lines, and multiple tissue samples from rodents treated *in vivo* (**Supporting Information Table 1**). In total, data from almost 1,000 individual treatments was reassessed, comprising 597 unique chemical entities including nucleobase analogues, commonly prescribed medications, bioactive molecules, and potential environmental toxins. An acknowledged limitation of this work is that the original choice of the agents was subjective (often a focus on toxicity) and the effects of individual drugs are often restricted to a single study, preventing a broader comparisons of drug effects between tissues/cell types. The cgr(base) quantification approach was applied to all 13 studies and each drug’s actions visualised by plotting cgr(C) against cgr(A). As an example, **Fig.3a** plots global gene expression profile changes in rat bone marrow after treatment with 76 drugs/agents (GSE59894, [4]). The diagonal bidirectionality is displayed by the majority of outlier drug effects, but some drugs show an asymmetric effect, perhaps indicating a more complex nucleobase deficit: composition interaction. Radar plots allow full visualisation of nucleobase influences from example drugs with symmetric (**Figs.3b C-A+ and 3c C+A-**) and asymmetric (**Fig.3d**) effects on global expression.

**Figure 3.**
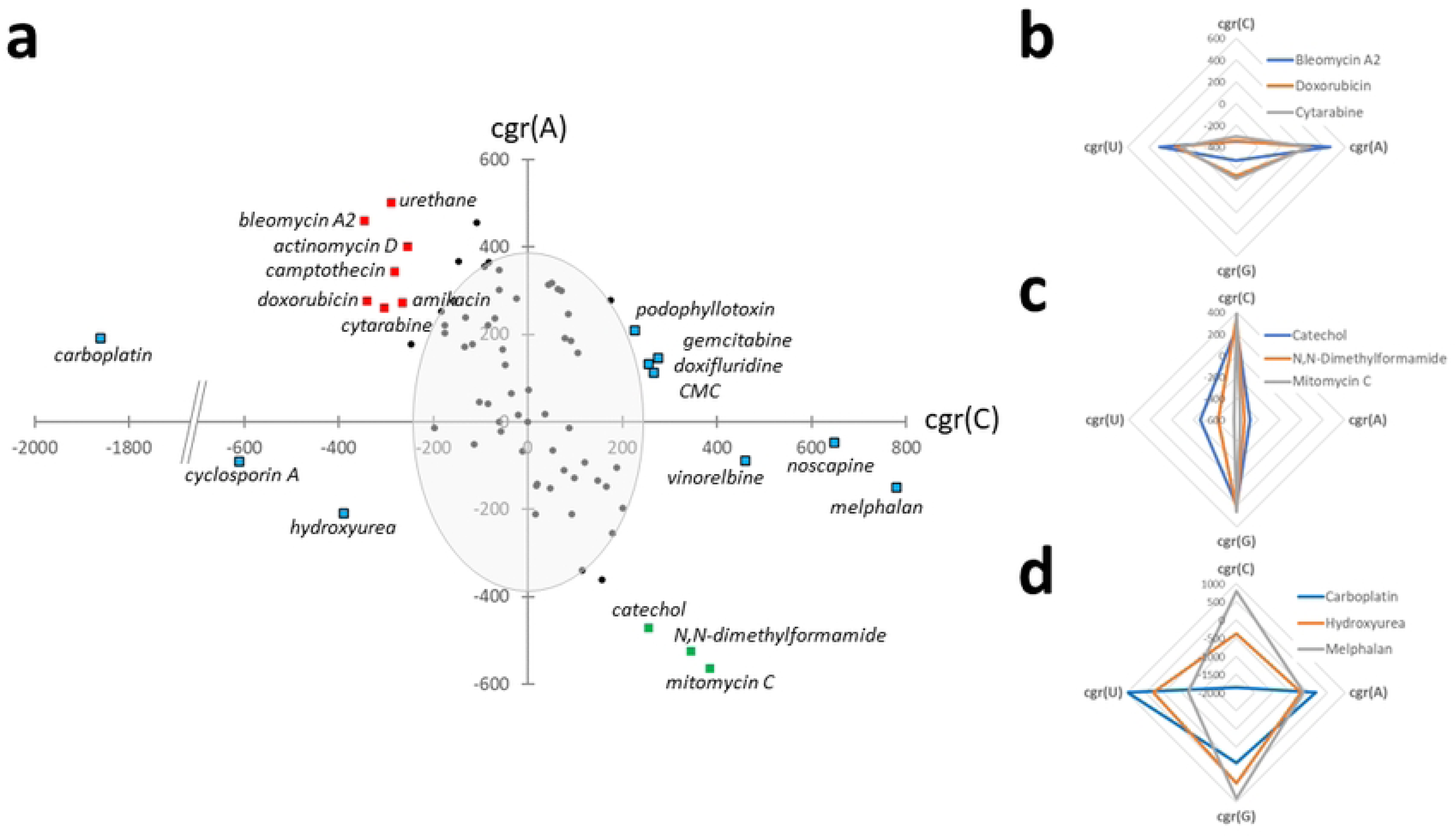
Representative study from the drug screen highlighting outlier molecules with substantial *in vivo* effects on the global transcriptional profile of rat bone marrow cells. Global gene expression profile changes in rat bone marrow after treatment with 76 drugs/agents (GSE59894, [4]). Data are plotted on a cgr(C) vs cgr(A) scatterplot where cgr represents the control corrected gradient of the expression profile for that base (a). The distribution is centred around the origin but extends primarily diagonally, top-left to bottom-right. Data points represent the average of approximately twelve replicate treatments. The control standard deviations for the two axes are indicated by the grey oval and, to be considered an outlier, the drug/chemical entity had to be located outside of this modest threshold. Green squares indicate outlier (C+A-) drugs inducing substantial expression increases in transcripts with higher proportion of C bases and concomitant decreases in those with increased A proportions. Red squares highlight drugs with outlier (C-A+) effects in the opposite direction. Although the majority of drug treatments caused symmetric profile changes along the diagonal C-A+ to C+A-axis, several drugs deviated from symmetry, as denoted by blue squares. Carboplatin has the moist extreme, assymmetric C-shift requiring the X-axis to be cropped. Radar plots allow the simultaneous visualisation of all four cgr(nucleobase) values for a drug, with the radius representing the effect size and direction (0 = control). Illustrative symmetric effect drugs (**Figs.3b C-A+ and 3c C+A-**), and asymmetric effect drugs (**Fig.3d**) cause characteristic global expression profile perturbations.

The published toxicities and historical market withdrawals identified for many extreme outlier drugs indicated that the magnitude of cgr(X) in any direction is probably limited by biological viability and adverse effects. Thus, to be categorised as an outlier, an agent was only required to exceed one standard deviation of the solvent control mean along at least two of the four nucleobase dimensions (approximately indicated by the grey oval in **Fig.3**). **Fig.4** lists the identities and clinical and molecular properties of the 131 treatments (115 unique entities, 19.3% of total) across the 13 studies that achieved such expression distortion, grouped by radar plot shape (symmetric C-A+ or C+A-directions, or asymmetric). Numerical data from all 13 studies are available in tabulated form in **Supporting Information Table 2**.

**Figure 4.**
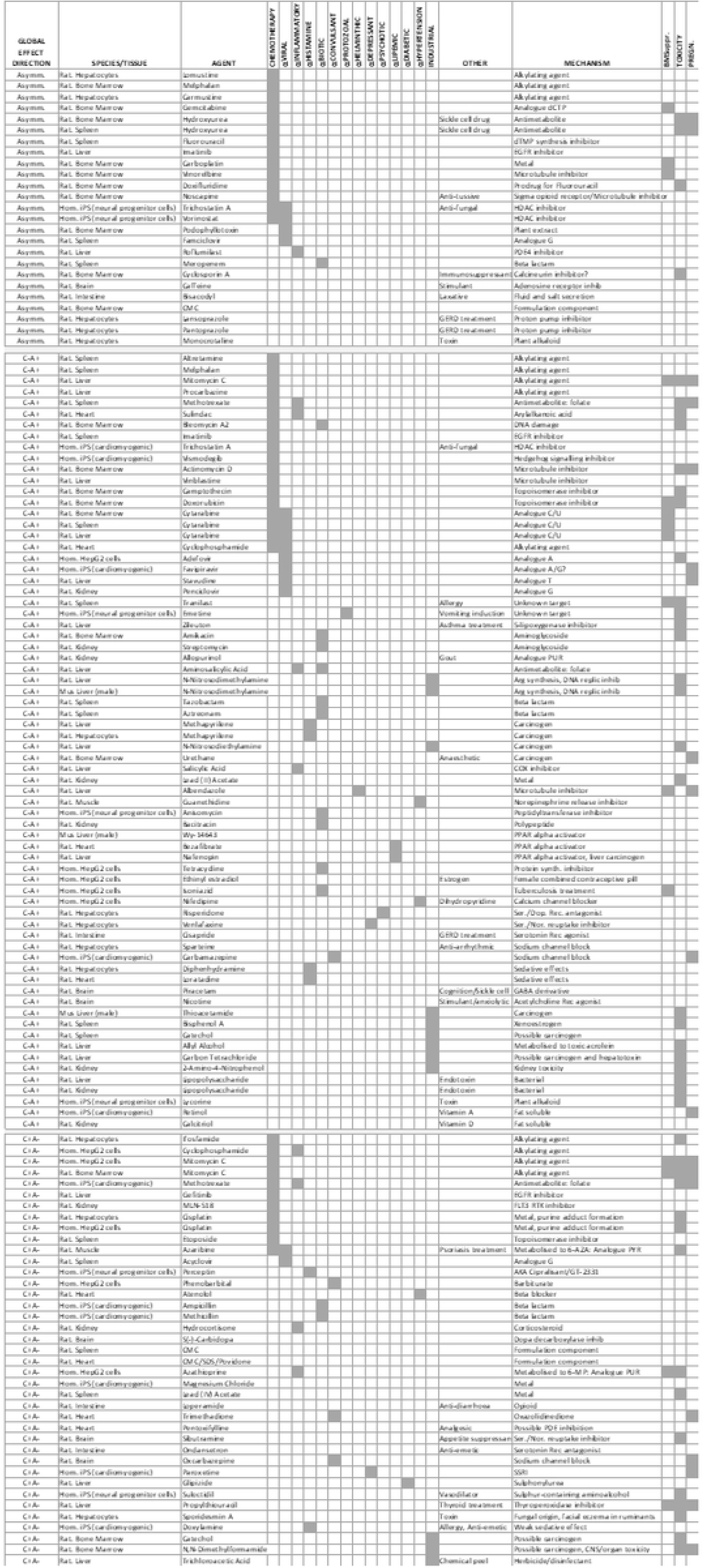
Chemical drug/entity treatments that induce a substantial change in global transcription expression profile. Entities are grouped according to direction of effect on profile, experimental tissue, details of therapeutic mode/mechanistic action/chemical nature, and reported side-effects. (Abbreviations: BMSuppr., bone marrow suppression; PREGN., known cause of teratogenicity or pregnancy risk; Ser., serotonin; Dop., dopamine; Nor., norepinephrine; PDE, phosphodiesterase; SSRI, selective serotonin reuptake inhibitor).

Importantly, nucleobase analogues and antimetabolite drugs are present within **Fig.4** indicating that a nucleobase supply mechanism is consistent with the global perturbations observed. Independent replications add to the robustness of the findings. The analogue cytarabine and alkylating agent mitomycin C were both outliers in three independent studies; and cisplatin, N-nitrosodimethylamine, catechol, CMC (carboxymethyl cellulose, a thickening agent used in food, cosmetics, and pharmaceuticals), methapyrilene, hydroxyurea, trichostatin A, imatinib, cyclophosphamide, lipopolysaccharide, melphalan, and methotrexate, were outliers in two independent studies. However, the direction of transcriptional distortion for specific drugs was occasionally different between tissues, such as in the case of methotrexate showing a C-A+ shift in the rat spleen but C+A-shift in human iPS cells (cardiomyogenic differentiation). Moreover, cisplatin has profound C+A-axis effects on expression in rat hepatocytes and human HepG2 cells, but little effect in rat kidney, bone marrow, or spleen. Tissue-/experiment-selective effects were observed for many other drugs perhaps reflecting differing *in vivo* absorption, distribution, metabolism, and excretion properties between tissues and cells.

Some chemical structure/function correlations were clear within the data. Analogue doxifluridine and its metabolite, fluorouracil, were independently identified as outliers, as were the epigenetic modifiers trichostatin A and chemical derivative, vorinostat. Gefitinib and imatinib, rather distant cousins within the EGFR inhibitor family, were both outliers but erlotinib (structurally very similar to gefitinib), was not. Proton pump inhibitors lansoprazole and pantoprazole were outliers. Related rabeprazole was similar in its direction of transcriptional effect, if not the magnitude required to be defined as an outlier, but a fourth member of the family, omeprazole, exhibited almost no effect on global transcription.

### Tissues show distinct baseline global transcriptional profiles, correlating with differentiation state

In the analysis above, tissues and cell types differed from each other in both the proportion of drug treatments that perturbed global expression profiles as well as the magnitude of those perturbations. The spleen showed the most robust outlier population, while kidney showed the least. This prompted an analysis of baseline tissue global expression profiles as a possible explanation. Three independent transcriptomic datasets of mouse organ/tissue/cell line expression profiles (GSE24207, GSE10246, GSE9954) were assessed (**Supporting Information Table 1**). All three produced largely similar gr(base) profiles, with GSE9954 shown in **Fig.5**. Among these, mouse spleen showed an extreme C-A+ distortion, perhaps rendering it more susceptible to transcriptional drug effects, whereas the kidney is more centrally located, perhaps making it more resistant to change. A key observation is that fetal tissues and ovary, together with embryonic stem, transformed, and haematopoetic cell lines occupy an extreme C-A+ region of the plot, suggesting that differentiation state or proliferative potential correlate with profile position on the bidirectional axis. The radar plot inset confirms the reduction in gr(C) in undifferentiated tissues, but also highlights a large increase in gr(G). Testis, despite its extraordinarily offset outlier position (due to its exceptionally negative gr(U)) also shares the large increase in gr(G).

**Figure 5.**
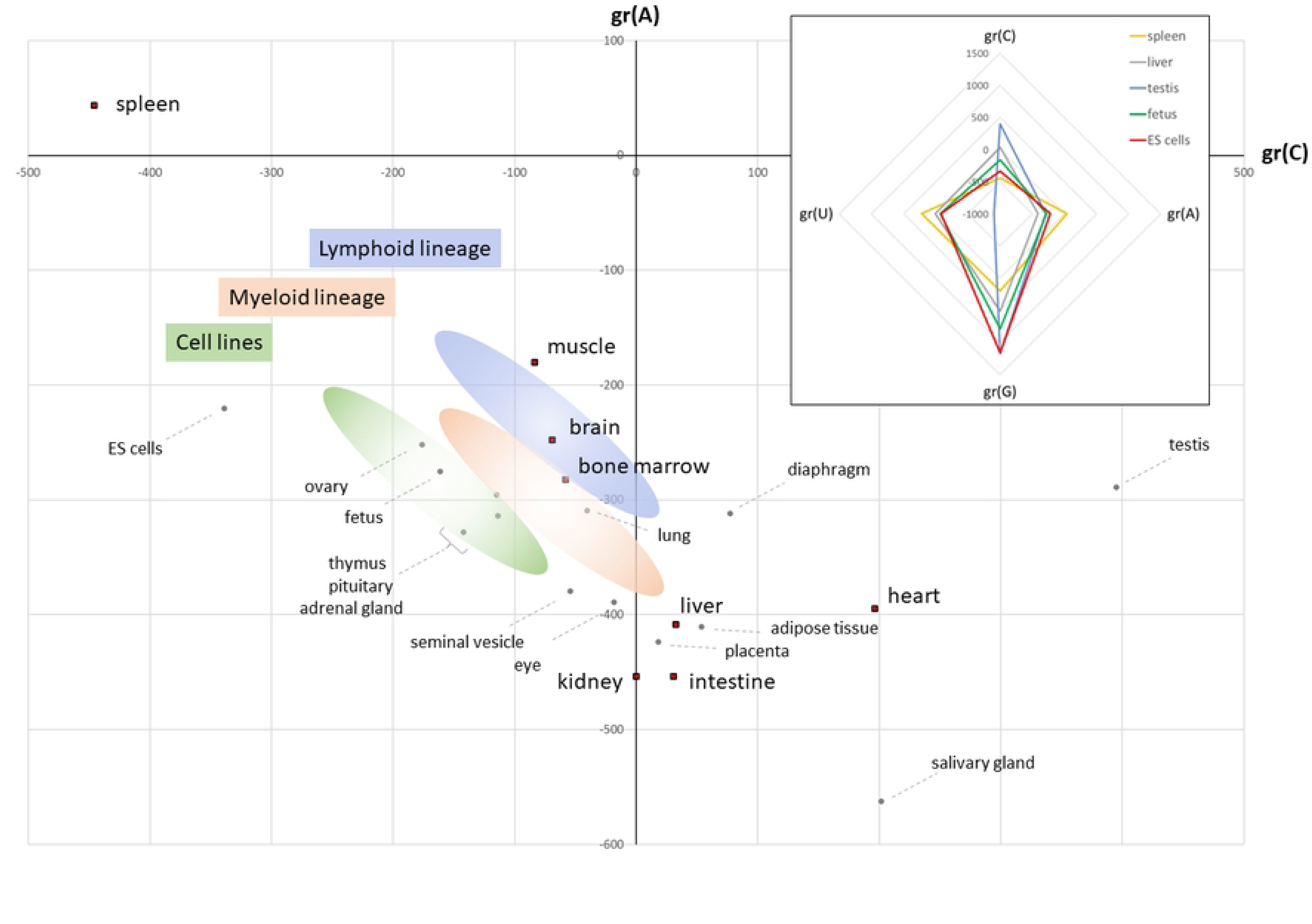
Organs and cell types possess distinct global transcriptional profiles that may contribute to their unique gene expression repertoire, drug sensitivity, and differentiation status. Tissue global expression profile from study GSE9954. Tissues employed in the drug screen are indicated by red squares, highlighting the link between the spleen’s extreme location and greatest perturbation by drugs. Coloured ovals indicate approximate positions for transformed cell lines and cells from the lymphoid and myeloid lineages, derived from GEO10246 data. The inset shows profile gradients for all four bases for four tissues and ES cells. The asymmetric location of testis in the main image is explained by the extreme negative gr(U) value in the radar plot.

### Physiological cycles, viral infections, a disorder of nucleobase supply, and multiple common, complex genetic disorders all significantly perturb global expression profiles

To determine if disease state or physiological change were associated with global expression profile shifts, gr(C) and gr(A) values were calculated from 30 publicly available array-based gene expression studies (**Supporting Information Table 1**, **Fig.6**).

**Figure 6.**
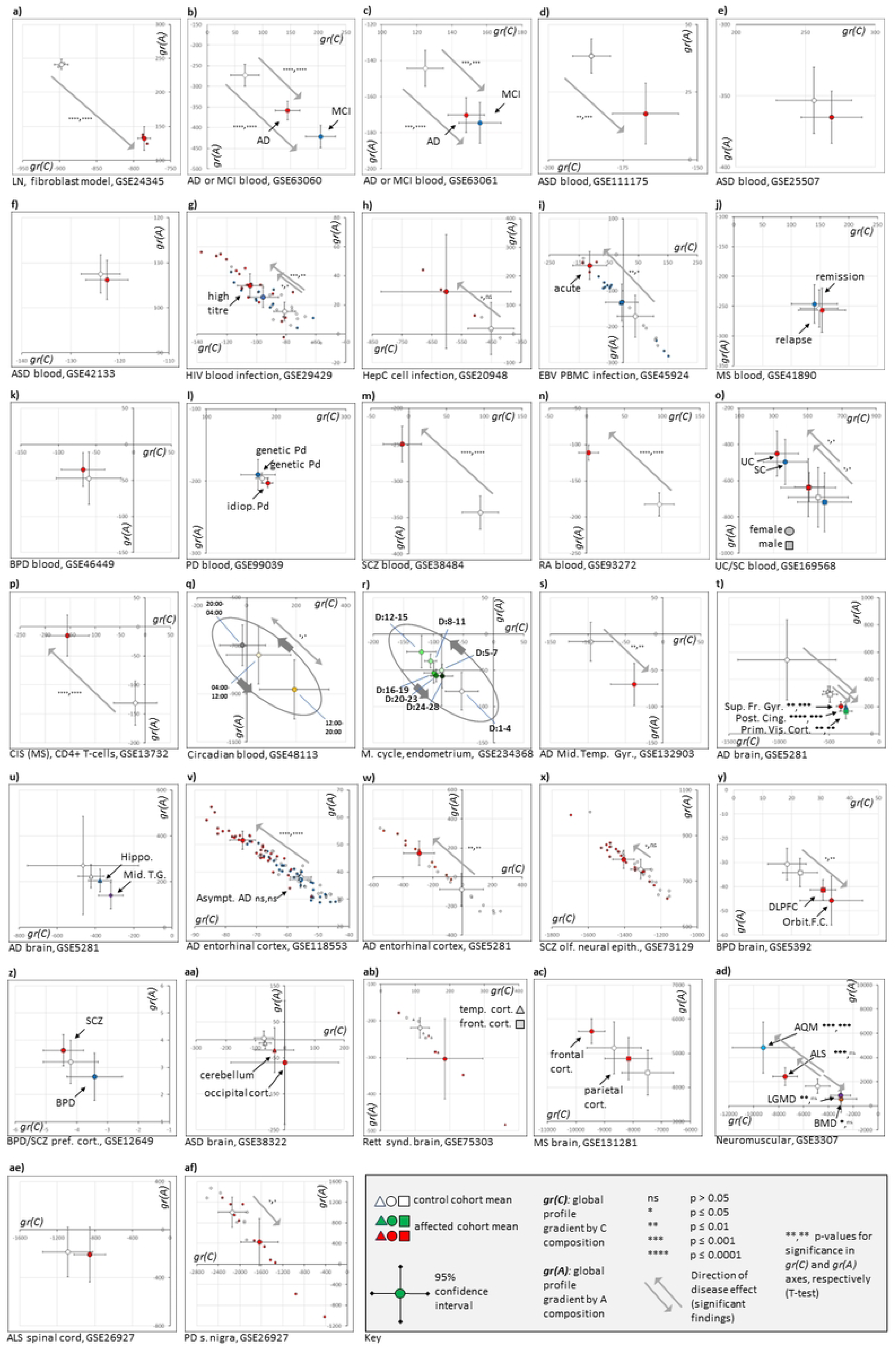
Viral infection and multiple complex genetic disorders significantly and bi-directionally perturb global transcriptomic profiles in blood, cells, and tissues. The panel of images plot gr(C) against gr(A) to represent the global transcriptional profile. Open shapes with error bars (95% confidence interval, conservative T-test statistic) represent mean values for uninfected or healthy control cohorts. Coloured shapes indicate the corresponding means for the comparative infected/disease state. Where relevant, statistically significant differences (two-tailed T-test) between healthy and disease gr(C) or gr(A) values, respectively, are indicated by asterisks, with arrows indicating effect direction. For viral response and some later tissue studies, where numbers of samples do not obscure interpretation, individual data are also shown alongside means to illustrate variance. The GEO public expression data accession (GSExxx) and brief description are found under each plot, but full descriptions of sample sizes and analysis are in Supporting Information Table 1. For 5q and 5r, oval shapes indicate the cyclical passage of time (24-hr clock or D: days) for circadian and menstrual rhythms. For Rett syndrome analysis (ab), the circles represent the averages and CIs of 6 pooled affected or 6 pooled healthy control samples. (Abbreviations - AD: Alzheimer’s disease (Asympt.: pathology but no cognitive effects), LN: Lesch-Nyhan syndrome, MCI: Mild cognitive impairment (prodromal AD), ASD: Autism spectrum disorder, HIV: Human immunodeficiency virus, HepC: Hepatitis C virus, EBV: Epstein-Barr virus, PBMC: Peripheral blood mononuclear cells, MS: Multiple sclerosis, BPD: Bipolar disorder, PD: Parkinson’s disease (idiopathic or genetic), SCZ: Schizophrenia, RA: Rheumatoid arthritis, UC: Ulcerative colitis, SC: A symptomatic form of inflammatory bowel disease which does not meet criteria for a diagnosis (e.g., of UC), CIS: Chronic isolation syndrome (prodromal MS), M. cycle: Menstrual cycle, Mid. Temp. Gyr.: Middle Temporal Gyrus of the brain, Sup. Fr. Gyr.: Superior frontal gyrus of the brain, Post. Cing.: Posterior cingulate cortex of the brain, Prim. Vis. Cort.: Primary visual cortex of the brain, Hippo.: Hippocampus of the brain, DLPC: Dorsolateral prefrontal cortex of the brain, Orbit. F. C.: Orbitofrontal cortex of the brain, Pref. cort.: Prefrontal cortex of the brain, Front. cort.: Frontal cortex of the brain, Temp. cort.: Temporal cortex of the brain, AQM: Acute quadriplegic myopathy, ALS: Amyotrophic lateral sclerosis (motor neuron disease), LGMD: Limb girdle muscular dystrophy, BMD: Becker muscular dystrophy, S. nigra: Substantia nigra of the brain).

Lesch-Nyhan (LN) syndrome, a purine salvage disorder caused by *HPRT* gene mutation, presents with complex neurological and neuropsychiatric symptoms [7]. *In vitro* cellular knockdown data (GSE24345, Fig.6a) [8], when reanalysed, demonstrated a highly significant C+A-global transcriptome change – a genetic counterpart to the earlier nucleobase analogue drug treatments.

In blood samples, a significant profile shift in the same, C+A-, direction as LN syndrome was observed in a pair of studies (GSE63060/63061, Fig.6b,c) of blood samples from those diagnosed with Alzheimer’s disease (AD) or mild cognitive impairment (MCI). The greater impact of MCI of profile may reflect early-stage pathology that becomes less detectable in full AD as neurons disappear. A similar shift was observed for one study (GSE111175, Fig.6d) of autism spectrum disorder (ASD), but two other ASD studies (GSE25507/42133, Fig.6e,f) failed to reach significance, even if their profiles trended in the same direction. Purine metabolism defects have been previously described for both AD [9] and ASD [10]. Together with observations of a shared transcriptional profile between AD and LN [11], these findings suggest commonalities in biochemical disturbance are reflected at the global transcriptional level.

Infection by human immunodeficiency virus (HIV, GSE29429, Fig.6g), Hepatitis C virus (HepC, GSE20948, Fig.6h), and Epstein-Barr virus (EBV, GSE45924, Fig.6i) produced significant shifts in the opposite, C-A+, direction. Dose-dependent effects were observed in samples with higher viral titre (>100,000) for HIV, and in the acute primary infection state for EBV. This suggests that early-stage, active virion production might drain the host cellular C/G nucleobase supply available for transcription, a process that that may align with other more recognised ‘host shutoff’ mechanisms [12].

No significant global transcriptional changes were seen in blood samples for multiple sclerosis (MS, GSE41890, Fig.6j), bipolar disorder (BPD, GSE46449, Fig.6k), or Parkinson’s disease (PD, GSE99039, Fig.6l). However, a significant C-A+ change for SCZ was observed in study GSE38485 (Fig.6m). Similarly, rheumatoid arthritis (RA, GSE93272, Fig.6n) presented with a significant C-A+ profile shift. Analysis of blood transcriptomics data from ulcerative colitis (UC) and symptomatic individuals not meeting UC diagnostic criteria (SC) (GSE169568, Fig.6o), showed trends in the C-A+ direction that were only statistically significant when females diagnosed with UC were compared to healthy control females, or when SC females were compared to SC males, indicating a potential influence of biological sex on phenotype. Crohn’s disease (not shown) did not show effects on global transcription. Clinically isolated syndrome (CIS), a precursor to full MS, produced a significant C-A+ global profile change in the CD4+T cell component of blood (GSE13732, Fig.6p). Finally, blood samples taken across the circadian day (GSE48113, Fig.6q) showed a C+A- shift during the afternoon/evening (12:00 to 20:00) and a C-A+ shift at night (20:00-0400). Mouse liver exhibits circadian fluxes in nucleobase biochemistry, especially purine metabolism, providing a necessary mechanistic link between circadian rhythm and the global expression changes observed [13]. Such shifts are likely to be a confounder for all the above transcriptomic studies where blood drawing was not at a consistent time of day. They may also provide an explanation for some aspects of chronotherapy where time of administration of drugs from Table 1 may synergise or antagonise with the circadian ebb and flow of changes in global expression profile [14–16].

Re-analysis of gene expression data from endometrial cell biopsies obtained across the menstrual cycle (GSE234368, Fig.6r) revealed another instance of cycling of the global gene expression profile: with the early follicular/proliferative phase (labels D1 through D15 indicating days post-menstruation) associated with a progressive C-A+ shift followed, after ovulation, by a C+A- return during the luteal/secretory phase (D16-D28). These changes may reflect the proliferation state of endometrial cells or the changing hormonal milieu (estrogen/FSH/LH/progesterone). The transcript encoding luteinizing hormone B chain (*LHB*) is in the top 0.5% of all transcripts for C-richness suggesting that its production in the pituitary gland may also be subject to global transcriptional control. Ethinyl estradiol is a component of the combined contraceptive pill and has a C-A+ activity in **Fig.4**. Similarly, bisphenol A, a toxic xenoestrogen identified with C-A+ action in the drug screen, is known to affect female reproductive health in numerous ways including menstrual irregularity and endometriosis [17].

In the assessment of other non-blood tissues in disease states, studies GSE36980 and GSE118553 provided no evidence for global expression changes associated with AD within the frontal cortex, temporal cortex, or hippocampus (data not shown). However, GSE132903 (Fig.6s) showed a significant C+A- shift for the middle temporal gyrus. GSE5281 (Fig.6t) showed significant C+A- changes within the visual cortex, the superior frontal gyrus, and the posterior cingulate cortex, with suggestive trends for the middle temporal gyrus and the hippocampus. Intriguingly, the AD global shift was statistically significant but reversed in direction (C-A+) within entorhinal cortex samples from two independent studies, GSE118553 (Fig.6v) and GSE5281 (Fig.6w). This brain region has unique characteristics that may correlate with it being one of the earliest locations for detectable AD pathology [18].

For SCZ, significant C-A+ changes were observed in biopsied olfactory neural epithelium (GSE73129, Fig.6x). For BPD, significant perturbations in the opposite direction to SCZ were observed for prefrontal cortex and orbitofrontal cortex (GSE5392, Fig.6y). However, no transcriptional changes were observed for either SCZ or BPD in another study of the prefrontal cortex (GSE12649, Fig.6z), although directionality of effect was maintained for both conditions.

For ASD, global profiles showed non-significant C+A- trends in the cerebellum and occipital cortex of a small study (GSE38322, **Fig.6aa**), and for the temporal cortex and frontal cortex in a study of the related condition, Rett syndrome (GSE75303, Fig.6ab). However, both of these studies included affected individuals with extreme outlier profile changes, suggesting a subgroup of cases with significant nucleobase effects. Such an ASD subgroup with nucleobase metabolism defects has been documented previously [19–21]. In the parietal cortex and frontal cortex of those diagnosed with MS, a similar direction-of-effect was observed, but this was not statistically significant compared to healthy controls (GSE131281, Fig.6a**c**).

Finally, a range of muscular/neuromuscular/neurodegenerative conditions were examined in small-scale studies GSE3307 (**Fig.6ad**) and GSE26927 (**Fig.6ae/af**). Amyotrophic lateral sclerosis (ALS/MND) and acute quadriplegic myopathy (AQM) showed significant C-A+ shifts in global profile of muscle biopsies, while Becker muscular dystrophy (BMD) and limb girdle muscular dystrophy (LGMD) showed significant changes in the opposite direction. The ALS findings were not replicated in spinal cord samples, but PD was associated with a significant C+A- shift in substantia nigra tissue gene expression profile, the anatomical origin of its neurodegenerative pathology.

### Changes in global transcriptional profile engage ‘hardwired’ biological responses and alter disease gene expression

Although tissues, drugs, and diseases have been described here in terms of their impact on global expression, ultimately it is expression change at the *individual* gene/protein level that produces functional consequences. It was hypothesised that specific mRNA transcripts might have evolved compositional extremes so that cells could initiate a ‘hardwired’ expression response to any fluctuations in nucleobase availability. Expression changes in the proteins encoded by such extreme mRNAs might act to restore supplies of nucleobases or may offer survival/selective advantages in adverse metabolic conditions. Such a system would manifest as enriched/over-represented gene ontology terms associated with those mRNAs at compositional extremes. The 110,962 human GENCODE v44 protein-coding transcripts were ranked by single nucleobase composition, or A+U or C+G paired nucleobase composition. Those transcripts found at the most extreme 1% ends of each ranking were assessed using ENRICHR, GOnet, and WEB Gestalt online gene ontology and functionality resources [22–24]. Many statistically significant gene ontology enrichments were identified including C- or (C+G)- rich transcripts over-represented in terms Regulation of DNA-templated Transcription (GO:0006355, adjusted p-values 1.3 x 10^-12^ and 1.3 X 10^-5^, respectively), and DNA-binding transcription factor activity, RNA polymerase II-specific (GO:0000981, FDR adjusted p-values 0.0050 and 1.7 x 10^-6^, respectively). Individually, C-richness was associated with Biological Process Involved in Intraspecies Interaction Between Organisms (GO:0051703, FDR adjusted p-value 0.0054) and G-richness with Regulation of Cellular Component Organization (GO:0051128, FDR adjusted p-value 1.8 x 10^-3^). It would be predicted that expression of genes with these GO terms would be diminished as global profiles shift in the C-A+ direction.

At the other extreme, (A+U)-rich transcripts were over-represented in Meiotic Cell Cycle (GO:0051321, FDR adjusted p-value 0.014) and Synaptonemal Complex Organization (GO:0070193, FDR adjusted p-value 0.015). Single nucleobase analysis of U-rich transcripts identified gene ontology terms Nervous System Process (GO:0050877, FDR adjusted p-value 0) and Detection of Chemical Stimulus (GO:0009593, FDR adjusted p-value 6.3 x 10^-28^) because of large clusters of olfactory and gustatory receptors at this extreme. A-rich transcripts, by contrast, were associated with Cilium Assembly (GO:0060271, FDR adjusted p-value 4.5 x 10^-5^), Cell Cycle (GO:0007049, FDR adjusted p-value 1.5 x 10^-9^), and Nuclear Chromosome Segregation (GO:0098813, FDR adjusted p-value 4.2 x 10^-8^). Genes with these GO terms would have reduced expression when global profiles shift in the C+A- direction.

Together, these data suggest that any biochemical stressor which compromises nucleobase availability will potentially restrict metabolic expenditure by affecting rates of cellular transcription or proliferation, while simultaneously engaging in behaviours which promote survival through greater resource-seeking or restricted resource allocation. This narrative closely resembles our previous analysis of protein functions at the extremes of nutritional amino acid composition [1].

Figures 7a-f describe the range of nucleobase compositions and explore the functionalities of extreme mRNAs. A frequency distribution of A+U composition in the entire human transcriptome is shown in **Fig.7a**. Aligned directly underneath are A+U vs. C composition scatterplots for all transcripts in grey, with the mean and +/- 2 standard deviations of A+U indicated (**Fig.7b**). The central 98% of transcripts are also indicated by the red bracket. Superimposed are key gene groupings selected for display due to their extreme compositions and established roles in important physiological processes and disease susceptibility.

**Figure 7.**
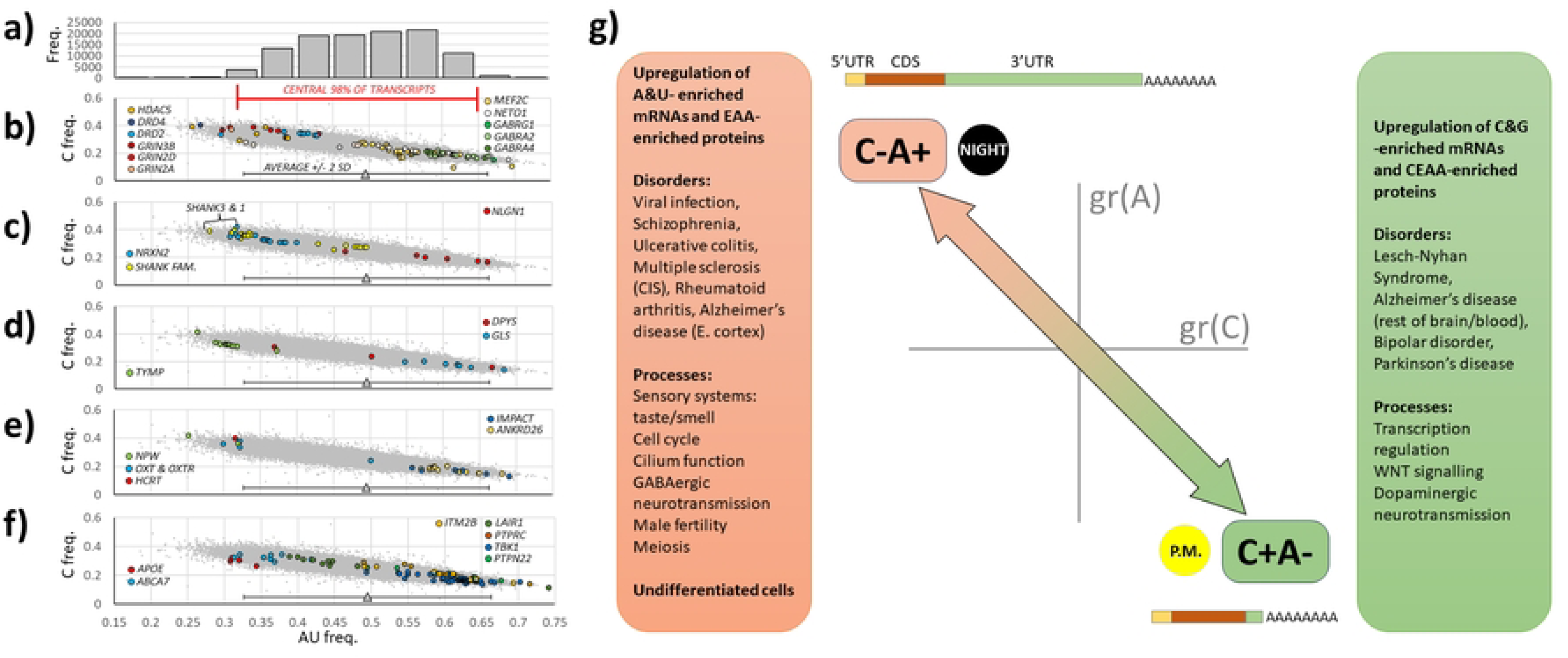
Many extreme composition mRNAs are disease susceptibility factors, contributing to the consequences of pathological global expression profile changes. The extremes of composition are illustrated by the histogram of transcripts within A+U frequency categories shown at the bottom (e.g., 0.45 indicates number of transcripts with A+U composition ≤ 0.45). The central 98% of all transcripts and +/- 2 standard deviation bars are presented to highlight extreme transcripts that fall outside of these limits. Extreme gene transcripts with key roles in brain function and disease (particularly neurodevelopmental disorders such as schizophrenia and autism spectrum disorder) are shown in b) and c). d) contains nucleobase metabolism genes which may be contributing to negative feedback control of shifts in global expression. In e), extreme transcripts dictating behavioural control of feeding are shown to occupy low A+U frequency extremes while the other end contains transcripts with cellular responses to nutritional deficit. f) contains major risk genes for Alzheimer’s disease and transcripts that may contribute to the immune/viral phenotypes associated with extreme C+A- and C-A+ shifts, respectively. The major directional shifts in global expression profile, C+A- and C-A+, are indicated on a summary representation of the gr(C) by gr(A) plot used throughout (g). These shifts occur over the circadian day (‘night’/’p.m.’) and are also associated with a diverging average transcript sizes and subregion proportions between the 500 most A+U rich mRNAs (2,891 nt), and the 500 most C+G rich (1,016 nt), as illustrated. The text boxes indicate the consequences of C+A- and C-A+ shifts on encoded protein composition, association with disorders, and over-represented gene ontology terms at those extremes.

The brain appears to use extreme composition to regulate development and activity (**Figs.7b & 7c**). The top 1% (C+G)-rich (=A+U-poor) transcripts include neurotransmitter receptors *GRIN3B*, *GRIN2D*, and *GRIN2A* (NMDA excitatory glutamatergic), *DRD2/4* (dopamine), and *CHRNA4* (acetylcholine); acetylcholinesterase, *ACHE*; neuronal channels *KCNQ2* and *CACNG6*; synaptic structural proteins *SHANK1&3*; neuroligin 2 post-synaptic protein, *NLGN2* (often associated with inhibitory neurons), and its pre-synaptic interactor neurexin 2, *NRXN2*. At the other extreme, (A+U)-rich transcripts include the neurotransmitter receptors *GABRA2*, *GABRA4*, and *GABRG1* (inhibitory, GABAergic); channels *SCN2A*, *KCNT2*, *CACNA2D* and *CACNB4*; neuroligin 1, *NLGN1* (often associated with excitatory neurons), and its interactor, neurexin 1, *NRXN1*. *HDAC5* is at the opposite extreme to the *MEF2C* gene it suppresses.

Together with extreme transcript, *NETO1* (a GRIN2A interactor), these three proteins are known to modulate synaptic plasticity. This catalogue of opposing distributions of opposing functionalities suggests that the brain can leverage nucleobase-mediated shifts in global transcriptional profile to control the tone of neuronal and circuit activity (excitatory vs. inhibitory) while also modulating trans-synaptic interactions. It also renders these genes and their functional systems susceptible to dysregulation by pathological changes in global profile.

Multiple nucleobase metabolism enzymes, such as TYMP and DPYS, are encode by transcripts located at compositional extremes (**Fig.7d**), potentially offering the hardwired feedback regulation by nucleobase availability suggested earlier. Two (A+U)-rich transcripts, *GLS* (glutaminase) and *ASNS* (asparagine synthetase), encode the enzymes that catabolise L- glutamine, the single amino acid starting-point for purine and pyrimidine base synthesis. A cell with reduced glutamine resources would have limited nucleobase synthesis, potentially lowering *GLS*/*ASNS* transcription levels and therefore preventing further glutamine loss through catabolism. Such a homeostatic mechanism has received experimental support from metabolomic analysis of *GLS* hypermorphs in which nucleobase synthesis is impaired [25]. Transcripts linking nucleobase deficits with homeostatic feeding behaviour outputs are found in the (C+G)-rich extremity (**Fig.7e**): *NPW* (feeding and drinking behaviours), oxytocin (*OXT*) and its receptor (*OXTR*)(social/offspring bonding, reproduction, and food consumption), and hypocretin (*HCRT*/orexin, arousal, hunger). At the A+U extreme, *IMPACT* and *ANKRD26* are two genes which mediate cellular adjustments to nutritional deficiency. Collectively, these nutritional findings directly complement our published protein analysis [1] in which EAA-rich proteins within the leptin signalling pathway provided a novel mechanistic connection between amino acid deficiency and hunger response.

Established individual neurodevelopmental, neurodegenerative, and autoimmune disease susceptibility genes such as *DRD2*, *APOE*, *ABCA7*, *TGFB1*, *TERT*, *POMC*, *HDAC5*, *WNT1*, *CARD9*, *LRP5*, *NRXN2*, *SHANK3&1*, *HBA*, *TH*, *PRRT2* and *IGF2* are rich in C+G, whereas *SNCA*, *PTPN22*, *LRRK2*, *TLR1*, and *PTH* are rich in A+U nucleobases. Their susceptibility to global expression change may help explain the disease findings presented in Fig.6 and may even contribute to the nosological classification of these disorders. For example, Alzheimer’s disease susceptibility genes *ABCA7* and *APOE* are strongly enriched in C+G nucleobases, as shown in **Fig.7f**. At the other extreme, out of the total of 110,962 transcripts examined, one transcript of *LAIR1* was ranked 2^nd^ most A+U enriched, in close proximity to transcripts for *PTPRC* (‘CD45’, 6^th^), *TBK1* (12^th^), and *PTPN22* (111^th^). These four genes encode key components of the immunological and antiviral responses. Their expression is likely to be increased following the C-A+ expression changes associated with viral infection and autoimmune status, as set out in Fig.6. Returning to Alzheimer’s disease, the transcript ranked 26^th^ most A+U enriched encodes the ITM2B (BRI2) protein, an inhibitor of amyloid precursor protein processing by secretases [26].

## Discussion

Re-examination of transcriptomic datasets reveals that drug treatment and disease pathology can cause coordinated distortions of cellular gene expression that are independent of promoter activity. These distortions appear to result from interactions of nucleobase availability and mRNA nucleobase composition. As such, mRNA transcription joins DNA replication [27, 28] and protein translation [1] to complete the trio of large-scale cellular polymerisation reactions for which substrate concentration is a regulatory influence.

This system of global regulation can be usefully compared to the principles of market economics. Each mRNA sequence defines the *demand*, the nucleobase availability is the *supply*, and the *price* is the biosynthetic cost of each transcript (assumed to be inversely proportional to its expression). Changes to supply through drug/disease action alter the market equilibrium and the cost of biosynthesis of each mRNA, ultimately guiding each transcript to a new equilibrium of expression level. The global profile, therefore, can be thought of as the totality of all expressed transcript equilibria. However, because it is shown that mRNA base composition and transcript length are covariates, it is possible to construct a rival mechanistic model that uses transcript length (for example, through drug/disease effects on transcriptional processivity) as the primary determinant of the observed global gene expression effects. In favour of the nucleobase supply model are the actions of nucleobase analogues, the asymmetric effects of some drugs, and the Lesch-Nyhan syndrome findings, with the coding and UTR sequences effectively operating as a combined ‘antenna’, detecting restricted nucleobase supply. A second interpretational issue is whether *positive* changes to expression gradients in the C-A+ or C+A- directions actually exist as a consequence increases in nucleobase availability. An alternative possibility, favoured here, is that all expression changes may be negative - the result of degrees of nucleobase insufficiency - with apparently positive effects an artifact of transcriptomics data normalisation procedures. Absolute gene expression quantification technologies will be needed to resolve this issue. Altered transcriptional competence is experienced across the composition spectrum which produces the breadth of global expression changes observed here that involve a large proportion of the transcriptome. Such wide-ranging changes are entirely relevant in terms of summed phenotypic *effects* (disease presentation/drug effect/drug adverse effect) but have probably confounded most attempts to determine root *causes* of disease through transcriptomics. A relevant example of this effect is the pervasive transcriptome-wide gene expression changes associated with *MYC* gene deregulation. These have defied explanation through straightforward models of transcription factor action [29, 30] but MYC’s established role in nucleotide synthesis regulation [31] might suggest global expression perturbation as an alternative explanation. MYC’s additional (or potentially entirely linked) roles as a driver of pluripotency and cancer also match observations here of a highly skewed baseline transcriptional profile in transformed and undifferentiated cell types such as ES cells.

Observed global changes in expression profile were largely limited to two directions, here termed C+A- and C-A+. Four consequences of this constrained directionality are immediately apparent.

Firstly, the precise phenotypic outcomes will vary from one organ to another as each tissue’s unique fingerprint of actively expressed genes presents a different substrate for the profile- defining conditions to act on. This may explain why diseases with an involvement of global expression profile dysregulation have tissue-specific pathologies.

Secondly, the bidirectional mechanism offers a more nuanced interpretation of conventional models of genetic susceptibility to disease or drug response because risk-carrying variants located within genes at the extremes of composition (e.g. *SHANK3* for autism spectrum disorder or *APOE* or *ABCA7* for Alzheimer’s disease) will be subject to reduced or increased impact (expressivity/penetrance) according to the direction of profile perturbation. As such, global expression control is a potential channel for environmental influence on genetic risk in instances where GxE effects operate.

Thirdly, the bidirectional effect offers a unifying model to explain the observed similarities and co-morbidities for certain disorders: for example, the established links between schizophrenia (C-A+) and autoimmune conditions (C-A+) [32], and how viral infection (C-A+) can be a trigger for both [33, 34]. These may indicate synergistic interactions due to a shared direction of action. Conditions such as epilepsy, that share many features with other neuropsychiatric disorders and possess risk genes with extreme composition, should be considered prime candidates for future study on this basis.

Lastly, a drug acting to drive the global expression profile in one direction (e.g., methotrexate, C+A-) might derive its beneficial effects as a ‘DMARD’ by reversing an opposite expression profile shift caused by a pathological condition (e.g., rheumatoid arthritis, C-A+). Employing the same logic, cancer chemotherapy agents such as methotrexate, sulindac, cytarabine, trichostatin A, cyclophosphamide, and azaribine detailed in Fig.4 seem to defy simple treatment categories – possessing antiviral, anti-inflammatory, and epigenetic actions. It is tempting to speculate that these shared actions are all a manifestation of their directional impacts on perturbed global transcriptomes.

The range of chemicals listed in Fig.4 offer new perspectives on drug action and adverse effects. The 12 antibiotics were an unexpected discovery given their traditionally accepted modes of action. Beta-lactams were found to act in all directions, but the remaining antibiotic classes only move profiles in a C-A+ direction. A recent multi-omic study of pathogen response to antibiotics identified purine/pyrimidine metabolism perturbation as a major target [35]. If this effect is shared with eukaryotic cells *in vivo*, antibiotic effectiveness or side-effects might be associated with global changes to both host and pathogen transcriptomes. Supporting this, accumulating evidence indicates that use of supplemental antibiotics in cell culture media can directly affect eukaryotic cell behaviour [36].

More generally, side-effects and disease risks are reported for many of the prescription drugs listed in Fig.4. The idiosyncratic and shared nature of adverse effects (bone marrow suppression, developmental/teratogenic changes, ototoxicity, fatigue/sleep disturbance, xerostomia, keratoconjunctivitis sicca, headache, fertility, and Stevens-Johnson syndrome) when compared to the huge chemical diversity of drugs is a paradox that may find resolution through the shared dysregulation of the pathways and processes found at the extremes of base composition. For example, it could be hypothesised that transcripts of deafness-associated genes, *MSRB3*, *ESPN*, and *WHRN*, all in possession of extreme A+U/C+G compositions, are likely to be sensitive to any drug-mediated global profile change, with ototoxicity a potential outcome. Another example of a long-term drug risk, which has received recent attention, is the purported link between the use of proton pump inhibitors such as lansoprazole and pantoprazole (both present in Fig.4) and an increased risk of Alzheimer’s disease [37], a condition that is shown here to be strongly associated with shifts in global transcriptional profile. If drug side-effects are indeed linked to global gene expression regulation, new small molecules may benefit from gene expression profile analysis to detect changes that could predict off-target risks.

The tight correlation between extreme A+U:C+G compositions within the 5’UTR, CDS, and 3’UTR of mRNAs and the nutritional amino acid composition of the resulting proteins was entirely unexpected because the allocation of codons to amino acids is considered an ancient ‘frozen accident’ that occurred millions of years before amino acids became categorisable by their nutritional source [38]. However, it might be postulated that a group of nine amino acids with codons highly biased in A+U nucleobase composition represented a ‘fault line’ in the genetic code that the animal kingdom took advantage of by outsourcing their metabolically expensive biosynthesis to organisms occupying lower trophic levels – those amino acids becoming ‘essential’ from that point forward. This seemingly irrational biological step was perhaps made feasible because global expression regulation could reduce harmful consequences of EAA dietary shortfall. Instances of poor nutrition, and the resulting compromised metabolic state impacting nucleobase supply, could act as a ‘canary in the coal mine’ that would pre-emptively reduce transcription of extreme A+U composition mRNAs encoding those proteins particularly rich in the nine EAAs. Indeed, it could be further speculated that UTR effects on protein composition indicate that natural selection maximised the sensitivity of nucleobase supply deficit detection by transcripts, ensuring limitations on the unsupportable translation of extreme EAA proteins. mRNAs that are part of important biological processes could have subsequently co-opted A+U/C+G composition to control their own regulation. Both this nutritional connection and the previously described drug/toxin effects provide a clear and quantifiable route for GxE interactions that are an important feature of complex phenotypes and illness.

In summary, it is now important to place global expression regulation within the hierarchy of disease aetiopathology – how it is connected to the root cause of disease and where it sits in relation to the other global pathologies such as epigenetic dysregulation, oxidative stress, ER stress, and chronic inflammation. In a similar vein, a vital future step is determining whether therapeutic manipulation of the global expression profile is a viable approach to treat disease.

## Methods

All analysis, calculations, graphing, and statistics were carried out in Microsoft Excel.

110,962 human mRNA transcripts (the protein-coding component of the GENCODE v44) were downloaded in FASTA format from (https://www.gencodegenes.org/). Download of mRATBN7.2 (NCBI) was used for the rat transcript analysis. Download of Gencode.vM33 was used for the mouse transcript analysis. Nucleobase frequencies and transcript lengths were calculated using the same approach detailed in [1]. Combination frequencies were also calculated, e.g., A+U.

Microarray-based gene expression datasets were downloaded from Gene Expression Omnibus (NCBI) [39]. All deposited data within GEO adhere to the ethical NCBI/GEO Human Subject Guidelines, where required. No human gene expression data can be traced back to an individual. GEO datasets were accessed between November 2023 and October 2024. =VLOOKUP was used to link microarray chip identifiers from gene expression data to the correct row of the transcript analysis. To examine the composition effect of a specific nucleobase on expression, the spreadsheet was sorted by one of C/A/G/U/AU/CG/AG/CU frequencies. Expression data for each sample/experiment across transcripts (typically deposited in log2 transformed form) were divided into 25 equal-sized bins from the lowest to the highest frequency of the searched nucleobase composition. Within each bin, expression values were averaged. To calculate profile gradients, gr(X), for each sample or experiment, the inverse log of the 2^nd^ bin was subtracted from the inverse log of the 24^th^ bin and then the result divided by the change in nucleobase composition between the corresponding bins. The extreme (1^st^/25^th^) bins were not used because they often showed greater variance. Corrected gradients cgr(X) were calculated by subtracting experimental control gradients from experimental sample gradients.

For the screen, drugs listed in Table 1 deviated from the mean of the controls by greater than 1 standard deviation of the control values in at least two of the four dimensions.

## Supporting information captions

### Supporting Information Table 1

The Gene Expression Omnibus accession codes for all microarray gene expression studies re- analysed in this manuscript are detailed within this table. The associated publications are referenced and information on the breakdown of sample numbers in case:control studies provided.

### Supporting Information Table 2

Thirteen datasets were analysed to identify drugs/chemical entities with pharmacological actions on global gene expression. In this table the full corrected gradient data (cgr(X)) are provided for all studies, together with colour highlighting to indicate drugs producing gradients of profile that substantially deviated from the control condition averages by at least 1 standard deviation.

## References

1. Thompson R, Pickard BS. The amino acid composition of a protein influences its expression. PLoS One. 2024;19(10):e0284234. Epub 20241014. doi: 10.1371/journal.pone.0284234. PubMed PMID: 39401228; PubMed Central PMCID: PMCPMC11472945.

2. Lane AN, Fan TW. Regulation of mammalian nucleotide metabolism and biosynthesis. Nucleic Acids Res. 2015;43(4):2466–85. Epub 20150127. doi: 10.1093/nar/gkv047. PubMed PMID: 25628363; PubMed Central PMCID: PMCPMC4344498.

3. Moffatt BA, Ashihara H. Purine and pyrimidine nucleotide synthesis and metabolism. Arabidopsis Book. 2002;1:e0018. Epub 20020404. doi: 10.1199/tab.0018. PubMed PMID: 22303196; PubMed Central PMCID: PMCPMC3243375.

4. Gusenleitner D, Auerbach SS, Melia T, Gomez HF, Sherr DH, Monti S. Genomic models of short-term exposure accurately predict long-term chemical carcinogenicity and identify putative mechanisms of action. PLoS One. 2014;9(7):e102579. Epub 20140724. doi: 10.1371/journal.pone.0102579. PubMed PMID: 25058030; PubMed Central PMCID: PMCPMC4109923.

5. Barrett T, Edgar R. Gene expression omnibus: microarray data storage, submission, retrieval, and analysis. Methods Enzymol. 2006;411:352–69. doi: 10.1016/S0076-6879(06)11019-8. PubMed PMID: 16939800; PubMed Central PMCID: PMCPMC1619900.

6. Barrett T, Wilhite SE, Ledoux P, Evangelista C, Kim IF, Tomashevsky M, et al. NCBI GEO: archive for functional genomics data sets--update. Nucleic Acids Res. 2013;41(Database issue):D991–5. Epub 20121127. doi: 10.1093/nar/gks1193. PubMed PMID: 23193258; PubMed Central PMCID: PMCPMC3531084.

7. Garcia-Gil M, Camici M, Allegrini S, Pesi R, Petrotto E, Tozzi MG. Emerging Role of Purine Metabolizing Enzymes in Brain Function and Tumors. Int J Mol Sci. 2018;19(11). Epub 20181114. doi: 10.3390/ijms19113598. PubMed PMID: 30441833; PubMed Central PMCID: PMCPMC6274932.

8. Kang TH, Guibinga GH, Jinnah HA, Friedmann T. HPRT deficiency coordinately dysregulates canonical Wnt and presenilin-1 signaling: a neuro-developmental regulatory role for a housekeeping gene? PLoS One. 2011;6(1):e16572. Epub 20110128. doi: 10.1371/journal.pone.0016572. PubMed PMID: 21305049; PubMed Central PMCID: PMCPMC3030599.

9. Ansoleaga B, Jove M, Schluter A, Garcia-Esparcia P, Moreno J, Pujol A, et al. Deregulation of purine metabolism in Alzheimer’s disease. Neurobiol Aging. 2015;36(1):68–80. Epub 20140808. doi: 10.1016/j.neurobiolaging.2014.08.004. PubMed PMID: 25311278.

10. Geryk J, Krsicka D, Vlckova M, Havlovicova M, Macek M, Jr., Kremlikova Pourova R. The Key Role of Purine Metabolism in the Folate-Dependent Phenotype of Autism Spectrum Disorders: An In Silico Analysis. Metabolites. 2020;10(5). Epub 20200506. doi: 10.3390/metabo10050184. PubMed PMID: 32384607; PubMed Central PMCID: PMCPMC7281253.

11. Kang TH, Friedmann T. Alzheimer’s disease shares gene expression aberrations with purinergic dysregulation of HPRT deficiency (Lesch-Nyhan disease). Neurosci Lett. 2015;590:35–9. Epub 20150127. doi: 10.1016/j.neulet.2015.01.042. PubMed PMID: 25636690.

12. Gaucherand L, Gaglia MM. The Role of Viral RNA Degrading Factors in Shutoff of Host Gene Expression. Annu Rev Virol. 2022;9(1):213–38. Epub 20220607. doi: 10.1146/annurev-virology-100120-012345. PubMed PMID: 35671567; PubMed Central PMCID: PMCPMC9530000.

13. Fustin JM, Doi M, Yamada H, Komatsu R, Shimba S, Okamura H. Rhythmic nucleotide synthesis in the liver: temporal segregation of metabolites. Cell Rep. 2012;1(4):341–9. Epub 20120420. doi: 10.1016/j.celrep.2012.03.001. PubMed PMID: 22832226.

14. Allada R, Bass J. Circadian Mechanisms in Medicine. N Engl J Med. 2021;384(6):550–61. doi: 10.1056/NEJMra1802337. PubMed PMID: 33567194; PubMed Central PMCID: PMCPMC8108270.

15. Kaur G, Phillips CL, Wong K, McLachlan AJ, Saini B. Timing of Administration: For Commonly-Prescribed Medicines in Australia. Pharmaceutics. 2016;8(2). Epub 20160415. doi: 10.3390/pharmaceutics8020013. PubMed PMID: 27092523; PubMed Central PMCID: PMCPMC4932476.

16. Zhang R, Lahens NF, Ballance HI, Hughes ME, Hogenesch JB. A circadian gene expression atlas in mammals: implications for biology and medicine. Proc Natl Acad Sci U S A. 2014;111(45):16219–24. Epub 20141027. doi: 10.1073/pnas.1408886111. PubMed PMID: 25349387; PubMed Central PMCID: PMCPMC4234565.

17. Kawa IA, Akbar M, Fatima Q, Mir SA, Jeelani H, Manzoor S, et al. Endocrine disrupting chemical Bisphenol A and its potential effects on female health. Diabetes Metab Syndr. 2021;15(3):803–11. Epub 20210331. doi: 10.1016/j.dsx.2021.03.031. PubMed PMID: 33839640.

18. Goettemoeller AM, Banks E, Kumar P, Olah VJ, McCann KE, South K, et al. Entorhinal cortex vulnerability to human APP expression promotes hyperexcitability and tau pathology. Nat Commun. 2024;15(1):7918. Epub 20240910. doi: 10.1038/s41467-024-52297-3. PubMed PMID: 39256379; PubMed Central PMCID: PMCPMC11387477.

19. Dai S, Lin J, Hou Y, Luo X, Shen Y, Ou J. Purine signaling pathway dysfunction in autism spectrum disorders: Evidence from multiple omics data. Front Mol Neurosci. 2023;16:1089871. Epub 20230203. doi: 10.3389/fnmol.2023.1089871. PubMed PMID: 36818658; PubMed Central PMCID: PMCPMC9935591.

20. Kurochkin I, Khrameeva E, Tkachev A, Stepanova V, Vanyushkina A, Stekolshchikova E, et al. Metabolome signature of autism in the human prefrontal cortex. Commun Biol. 2019;2:234. Epub 20190621. doi: 10.1038/s42003-019-0485-4. PubMed PMID: 31263778; PubMed Central PMCID: PMCPMC6588695.

21. Zigman T, Petkovic Ramadza D, Simic G, Baric I. Inborn Errors of Metabolism Associated With Autism Spectrum Disorders: Approaches to Intervention. Front Neurosci. 2021;15:673600. Epub 20210528. doi: 10.3389/fnins.2021.673600. PubMed PMID: 34121999; PubMed Central PMCID: PMCPMC8193223.

22. Chen EY, Tan CM, Kou Y, Duan Q, Wang Z, Meirelles GV, et al. Enrichr: interactive and collaborative HTML5 gene list enrichment analysis tool. BMC Bioinformatics. 2013;14:128. Epub 20130415. doi: 10.1186/1471-2105-14-128. PubMed PMID: 23586463; PubMed Central PMCID: PMCPMC3637064.

23. Elizarraras JM, Liao Y, Shi Z, Zhu Q, Pico AR, Zhang B. WebGestalt 2024: faster gene set analysis and new support for metabolomics and multi-omics. Nucleic Acids Res. 2024;52(W1):W415–W21. doi: 10.1093/nar/gkae456. PubMed PMID: 38808672; PubMed Central PMCID: PMCPMC11223849.

24. Pomaznoy M, Ha B, Peters B. GOnet: a tool for interactive Gene Ontology analysis. BMC Bioinformatics. 2018;19(1):470. Epub 20181207. doi: 10.1186/s12859-018-2533-3. PubMed PMID: 30526489; PubMed Central PMCID: PMCPMC6286514.

25. Rumping L, Pras-Raves ML, Gerrits J, Tang YF, Willemsen MA, Houwen RHJ, et al. Metabolic fingerprinting reveals extensive consequences of GLS hyperactivity. Biochim Biophys Acta Gen Subj. 2020;1864(3):129484. Epub 20191114. doi: 10.1016/j.bbagen.2019.129484. PubMed PMID: 31734463.

26. Matsuda S, Senda T. BRI2 as an anti-Alzheimer gene. Med Mol Morphol. 2019;52(1):1–7. Epub 20180423. doi: 10.1007/s00795-018-0191-1. PubMed PMID: 29687167.

27. Bester AC, Roniger M, Oren YS, Im MM, Sarni D, Chaoat M, et al. Nucleotide deficiency promotes genomic instability in early stages of cancer development. Cell. 2011;145(3):435–46. doi: 10.1016/j.cell.2011.03.044. PubMed PMID: 21529715; PubMed Central PMCID: PMCPMC3740329.

28. Hastak K, Paul RK, Agarwal MK, Thakur VS, Amin AR, Agrawal S, et al. DNA synthesis from unbalanced nucleotide pools causes limited DNA damage that triggers ATR-CHK1-dependent p53 activation. Proc Natl Acad Sci U S A. 2008;105(17):6314–9. Epub 20080423. doi: 10.1073/pnas.0802080105. PubMed PMID: 18434539; PubMed Central PMCID: PMCPMC2359797.

29. Cotterman R, Jin VX, Krig SR, Lemen JM, Wey A, Farnham PJ, et al. N-Myc regulates a widespread euchromatic program in the human genome partially independent of its role as a classical transcription factor. Cancer Res. 2008;68(23):9654–62. doi: 10.1158/0008-5472.CAN-08-1961. PubMed PMID: 19047142; PubMed Central PMCID: PMCPMC2637654.

30. Nie Z, Hu G, Wei G, Cui K, Yamane A, Resch W, et al. c-Myc is a universal amplifier of expressed genes in lymphocytes and embryonic stem cells. Cell. 2012;151(1):68–79. doi: 10.1016/j.cell.2012.08.033. PubMed PMID: 23021216; PubMed Central PMCID: PMCPMC3471363.

31. Villa E, Ali ES, Sahu U, Ben-Sahra I. Cancer Cells Tune the Signaling Pathways to Empower de Novo Synthesis of Nucleotides. Cancers (Basel). 2019;11(5). Epub 20190517. doi: 10.3390/cancers11050688. PubMed PMID: 31108873; PubMed Central PMCID: PMCPMC6562601.

32. Cullen AE, Holmes S, Pollak TA, Blackman G, Joyce DW, Kempton MJ, et al. Associations Between Non-neurological Autoimmune Disorders and Psychosis: A Meta-analysis. Biol Psychiatry. 2019;85(1):35–48. Epub 20180628. doi: 10.1016/j.biopsych.2018.06.016. PubMed PMID: 30122288; PubMed Central PMCID: PMCPMC6269125.

33. Kotsiri I, Resta P, Spyrantis A, Panotopoulos C, Chaniotis D, Beloukas A, et al. Viral Infections and Schizophrenia: A Comprehensive Review. Viruses. 2023;15(6). Epub 20230609. doi: 10.3390/v15061345. PubMed PMID: 37376644; PubMed Central PMCID: PMCPMC10302918.

34. Sundaresan B, Shirafkan F, Ripperger K, Rattay K. The Role of Viral Infections in the Onset of Autoimmune Diseases. Viruses. 2023;15(3). Epub 20230318. doi: 10.3390/v15030782. PubMed PMID: 36992490; PubMed Central PMCID: PMCPMC10051805.

35. Yang JH, Wright SN, Hamblin M, McCloskey D, Alcantar MA, Schrubbers L, et al. A White-Box Machine Learning Approach for Revealing Antibiotic Mechanisms of Action. Cell. 2019;177(6):1649–61 e9. Epub 20190509. doi: 10.1016/j.cell.2019.04.016. PubMed PMID: 31080069; PubMed Central PMCID: PMCPMC6545570.

36. Ryu AH, Eckalbar WL, Kreimer A, Yosef N, Ahituv N. Use antibiotics in cell culture with caution: genome-wide identification of antibiotic-induced changes in gene expression and regulation. Sci Rep. 2017;7(1):7533. Epub 20170808. doi: 10.1038/s41598-017-07757-w. PubMed PMID: 28790348; PubMed Central PMCID: PMCPMC5548911.

37. Pourhadi N, Janbek J, Jensen-Dahm C, Gasse C, Laursen TM, Waldemar G. Proton pump inhibitors and dementia: A nationwide population-based study. Alzheimers Dement. 2024;20(2):837–45. Epub 20231005. doi: 10.1002/alz.13477. PubMed PMID: 37795826; PubMed Central PMCID: PMCPMC10917029.

38. Crick FH. The origin of the genetic code. J Mol Biol. 1968;38(3):367–79. doi: 10.1016/0022-2836(68)90392-6. PubMed PMID: 4887876.

39. Edgar R, Domrachev M, Lash AE. Gene Expression Omnibus: NCBI gene expression and hybridization array data repository. Nucleic Acids Res. 2002;30(1):207–10. doi: 10.1093/nar/30.1.207. PubMed PMID: 11752295; PubMed Central PMCID: PMCPMC99122.

